# SHAPE directed RNA folding

**DOI:** 10.1101/015537

**Authors:** Dominik Luntzer, Ronny Lorenz, Ivo L. Hofacker, Peter F. Stadler, Michael T. Wolfinger

## Abstract

**Summary:** Chemical mapping experiments allow for nucleotide resolution assessment of RNA structure. We demonstrate that different strategies of integrating probing data with thermodynamics-based RNA secondary structure prediction algorithms can be implemented by means of soft constraints. This amounts to incorporating suitable pseudo-energies into the standard energy model for RNA secondary structures. As a showcase application for this new feature of the `viennaRNA Package` we compare three distinct, previously published strategies to utilize SHAPE reactivities for structure prediction. The new tool is benchmarked on a set of RNAs with known reference structure.

**Availability and implementation:** The capability for SHAPE directed RNA folding is part of the upcoming release of the `viennaRNA Package 2.2`, for which a preliminary release is already freely available at http://www.tbi.univie.ac.at/RNA.

**Supplementary information:** Supplementary data is attached.

## 1 INTRODUCTION

Beyond its role as information carrier from genome to proteome, RNA is a key player in genome regulation and contributes to a wide variety of cellular tasks. The spatial structure RNA plays an important role in this context because it critically influences the interaction of RNAs with proteins and with nucleic acids. Knowledge of RNA structure is therefore crucial for understanding various biological processes. Chemical and enzymatic probing methods provide information concerning the flexibility and accessibility at nucleotide resolution. They are based on the observation that RNA can be selectively modified by small organic molecules, metal ions or RNAse enzymes, resulting in formation of an adduct between the RNA and the small compound or RNA cleavage. Subsequent primer extension mediated by RT enzymes typically terminates at the modified sites. The resulting cDNA fragments thus inform directly on the RNA structure by identifying, depending on the particular reagent, paired or unpaired sequence positions.

The first chemical probing workflows were developed in the 1980ies (Peattie and Gilbert, 1980; Stern *et al*., 1988), followed by novel approaches in the last 10 years, including protocols based on hydroxyl radicals (Tullius and Greenbaum, 2005), inline probing (Regulski and Breaker, 2008), dimethyl sulfate (DMS) (Cordero *et al*., 2012), and *selective 2’-hydroxyl acylation analyzed by primer extension (SHAPE)* (Merino *et al*., 2005; Weeks, 2010), which have recently been applied *in vivo* (Zemora and Waldsich, 2010; Wildauer *et al*., 2014; Kwok *et al*., 2013). Structural characterization of multiple RNAs in high throughput sequencing has been reported in different assays, including FragSeq (Underwood *et al*., 2010), PARS (Kertesz *et al*., 2010), SHAPE-seq (Lucks *et al*., 2011), and Mod-seq (Talkish *et al*., 2014). Moreover, a model-based approach for *in silico* assessment and characterization of large-scale NGS structure mapping approaches has recently been reported (Aviran and Pachter, 2014).

As chemical probing is becoming a frequently used technology to determining RNA structure experimentally, there is increasing demand for efficient and accurate computational methods to incorporate probing data into secondary structure prediction tools. Efficient dynamic programming algorithms, as implemented in the `ViennaRNA Package` (Hofacker *et al*., 1994; Lorenz *et al*., 2011), rely on the standard energy model for RNA folding, which is based on the assumption that the free energy of a given structure is composed of the free energies of its loops (Xia *et al*., 1998). While such a thermodynamics-based approach yields excellent prediction results for short sequences, accuracy decreases to between 40% and 70% for long RNA sequences. This discrepancy is mainly caused by imperfect thermodynamic parameters and the inherent limitations of the secondary structure model, such as tertiary interactions, pseudoknots, ligand binding, or kinetics traps. To alleviate the gap in available computational tools we have developed a framework for incorporating probing data into the structure prediction algorithms of the `ViennaRNA Package` by means of *soft constraints* in order to improve prediction quality.

## 2 METHODS

Compared to the application of *hard constraints* (Mathews *et al*., 2004), which restrict the folding space by forcing certain nucleotides to either being paired or unpaired, the incorporation of *soft constraints* is a less stringent approach. Soft constraints guide the folding process by adding position specific or motif specific pseudo-energy contributions to the experimentally determined free energy values of specific secondary structure motifs. This amounts to a distortion of the equilibrium ensemble of structure in favour of structures that are consistent with experimental data. Positive energy contributions are used to penalize mismatching motifs, while negative energy contributions give a “bonus” to those features that agree between prediction and experimentally determined pairing pattern.

The first approach that applied SHAPE directed RNA folding uses the simple linear ansatz

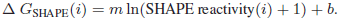

to convert SHAPE reactivity values to pseudo energies whenever a nucleotide *i* contributes to a stacked pair (Deigan *et al*., 2009). A positive slope *m* penalizes high reactivities in paired regions, while a negative intercept *b* results in a confirmatory “bonus” free energy for correctly predicted base pairs. Since the energy evaluation of a base pair stack involves two pairs, the pseudo energies are added for all four contributing nucleotides. Consequently, the energy term is applied twice for pairs inside a helix and only once for pairs adjacent to other structures. For all other loop types the energy model remains unchanged even when the experimental data highly disagrees with a certain motif.

A more consistent model considers nucleotide-wise experimental data in all loop energy evaluations (Zarringhalam *et al*., 2012). First, the observed SHAPE reactivity of nucleotide *i* is converted into the probability *q_i_* that position *i* is unpaired by means of a non-linear map. Then pseudo-energies of the form

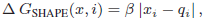

are computed, where *x_i_* = 0 if position *i* is unpaired and *x_i_* = 1 if *i* is paired in a given secondary structure. The parameter *β* serves as scaling factor. The magnitude of discrepancy between prediction and experimental observation is represented by |*x_i_ – q_i_*|.

These two methods incorporate pseudo-energies even when the observed data are consisted with an unaided secondary structure prediction. In a different approach, Washietl *et al*. (2012) suggested to phrase the choice of the bonus energies as an optimization problem aiming to find a perturbation vector 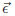
in such a away that the discrepancy between the observed and predicted probabilities to see position *i* unpaired, respectively, is minimized. At the same time, the perturbation should be as small as possible. The tradeoff between the two goals is naturally defined by the relative uncertainties inherent in the SHAPE measurements and the energy model, respectively. An appropriate error perturbation vector thus satisfies

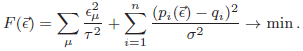

Here, *є_μ_* is the perturbation energy for a certain structural element *μ* and the variances 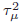 and 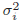 serve as weighting factors for the relative influence of the structure predicted from the standard energy model compared to the experimental data.

In this setting, the energy model is only adjusted when necessary. If the thermodynamic model already yields a perfect prediction, the resulting perturbation vector vanishes and the folding recursions remain unbiased. Otherwise the perturbation vector is used to guide the folding process by adding a pseudo-energy *є_i_* whenever nucleotide *i* appears unpaired in the folding recursions. Since the pairing probabilities are derived from the whole ensemble of secondary structures, the algorithm of Washietl *et al*. (2012) tends to decrease structural diversity only slightly, which makes it applicable to RNAs with several distinct low free energy structures. Furthermore, the inferred perturbation energies identify sequence positions that require major adjustments of the energy model to conform with the experimental data. High perturbation energies for just a few nucleotides are therefore indicative of posttranscriptional modifications or intermolecular interactions that are not explicitly handled by the energy model. A major drawback of this approach is its asymptotic time complexity of *O*(*n*^4^), which renders it very expensive for long sequences. This can be alleviated by a sampling strategy for estimating the gradient of the error functional *F* and provides a viable alternative to the exact numeric solution that reduces the time complexity to *O*(*n*^3^) again.

### Implementation

All three methods outlined above have been implemented into the `ViennaRNA Package`. Additional functionalities are available through the API of the `ViennaRNA Library` and through the command line interface of the application `RNAfold`. The required changes to the folding recursions and technical details of handling both hard and soft constraints in `ViennaRNA` will be described elsewhere in full detail. The key feature for our purposes is the consistent incorporation of a user defined position dependent energy contribution for each nucleotide that remains unpaired. The novel standalone tool `RNApvmin` dynamically estimated this vector of pseudo-energies that minimize model adjustments and discrepancies between observed and predicted pairing probabilities. Once this perturbation vector has been calculated, the pseudo-energies in the vector can be used to constrain folding with `RNAfold`. This setup makes is easy for users to incorporate alternative ways of computing bonus energies, e.g. along the lines of Eddy (2014), or to use the software with other types of probing data. The additional strategies for probing data/bonus energy incorporation into the folding recursions introduce a variety of parameters. These need to be chosen carefully. We refer to the supplement material for a detailed summary of the default parameters. The new features have been included into the `ViennaRNA Websuite` (Gruber *et al*., 2008), a Web server which provides many functionalities of the `ViennaRNA Package` and is available at http://rna.tbi.univie.ac.at.

## 3 RESULTS

We applied the methods to a benchmark set with known reference structures (Hajdin *et al*., 2013). The test set contains 24 triples of sequences, their corresponding SHAPE data, and reference structures. The reference structures were derived from X-ray crystallography experiments or predicted by comparative sequence analysis.

The use of soft constraints derived from SHAPE data leads to improved prediction results for many RNAs. This is clearly visible in the predictions from our benchmark data set (see Figure 1, and Supplementary data). However, for some of the RNAs within our benchmark data the additional pseudo-energy terms lead to worse predictions. This may be due to two factors. First, experimental data always comes with a certain inaccuracy. Second, the underlying energy model excludes pseudoknotted structures, which are present in approximately half of the benchmarked RNAs. Hence, pseudoknot interactions are not only present in the reference structure, but also influence the SHAPE reactivities.

**Fig. 1.**
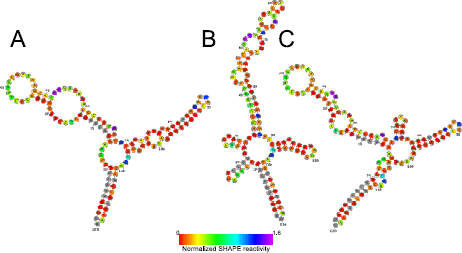
Secondary structure prediction of E.coli 5S rRNA from our benchmark data set. **A** Structure reference, **B** prediction by `RNAfold` with default parameters, and **C** prediction by `RNAfold` with guiding pseudo-energies obtained from SHAPE reactivity data using `RNApvmin`. Structure plots created using the `forna` webserver (Kerpedjiev and Hammer, 2015). Grey nucleotides correspond to missing SHAPE reactivity data.

Incorporation of probing data not only affects the minimum free energy structure, but also the entire ensemble of structures. Consequently, the predicted pairing probabilities are shifted towards the observed reactivity pattern. However, the effect is less distinct when applying the method of Washietl *et al*. (see Supplementary Figure 11). The incorporation of SHAPE reactivities into the folding recursion via soft constraints produces prediction results closer to the experimentally observed reference structures. However, none of the three approaches consistently outperforms the other two methods in terms of prediction sensitivity.

As an additional example, we compare the three methods using sequence and probing data of an artificially designed theophylline sensing riboswitch (Qi *et al*., 2012), see Supplementary data 4. The generalized handling of soft constraints in the `ViennaRNA Package` as of version 2.2 enables to directly include the ligand binding free energy of theophylline to the aptamer.

## ACKNOWLEDGEMENTS

This work was partly funded by the Austrian Science Fund (FWF) project “RNA regulation of the transcriptome” (F43), Deutsche Forschungsgemeinschaft (DFG) STA 850/15-1 and the German ministry of science (0316165C as part of the e:Bio initiative).

## Supplementary Material

### 1 Availability

The performed benchmark can be downloaded at http://github.com/dluntzer/
shapebenchmark and may be of further use when a novel method should be
compared against the existing approaches.

### 2 Metrics

#### 2.1 Minimum free energy structure

In order to rate the quality of RNA secondary structure prediction results in terms of prediction accuracy, the predicted minimum free energy structures are usually compared to known reference secondary structures. Suboptimal folds, yielding a free energy within a certain range from the minimum free energy structure, may also contain structures with similar or even better quality than the MFE structure. However, there is no way to rate the quality of suboptimal folds when the reference structure is unknown, which is usually the case. As a result suboptimal folds can not be used to rate the quality of a secondary structure prediction algorithm.

The correctness of the minimum free energy structure prediction is rated by comparing the predicted base pairs against the base pairs determined from the reference structure. In order to rate the quality, two parameters are commonly used to describe the amount of correct and wrong base pair predictions. The *Sensitivity* represents the percentage of base pairs in the reference structure, which are also found in the prediction. However, many RNA secondary structure prediction algorithms tend to predict additional pairs, which can not be verified with experimental methods. As a result the *Positive predictive value* (*PPV*), which represents the percentage of predicted pairs found in the reference structure, is used to rate the amount of false positives.

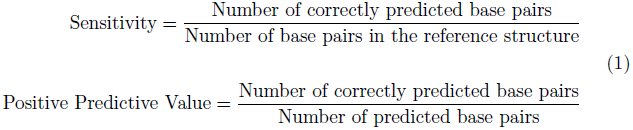

#### 2.2 Partition function

In contrast to the MFE structure, the partition function approach is used to model the whole ensemble rather than just predicting one or a few promising secondary structure candidates. Nevertheless, various parameters can be used to describe the agreement of the predicted ensemble with the known reference structure.

The *Pairing Probability Score* is defined as the arithmetic mean of the predicted pairing probabilities *p_ij_* of all pairs contributing to the reference structure *S* and shows the agreement of the pairing probability matrix derived from the ensemble of all possible structures with one single reference structure.

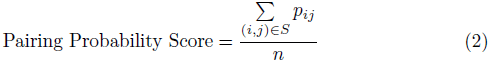

The *Ensemble Diversity* 〈*d*〉 shows the mean distance between predicted pairs, which can be obtained from the predicted pairing probabilities. Since the algorithms for incorporating experimental data tend to favor motifs that are in agreement with the observed experimental data, while penalizing disagreeing motifs, the *Ensemble Diversity* is used to illustrate to which extent the shift towards the experimental data influences the variability of the secondary structures represented by the ensemble. The *Ensemble Diversity* of a thermodynamic based prediction depends on the energy model and its parameters. Since the incorporation of additional experimental information is usually done by adding additional constraints, a decrease in the *Ensemble Diversity* in contrast to the thermodynamic based prediction is expected. Since the amount of possible pairs raises with growing sequence lengths, the *Ensemble Diversity* is normalized through division by the length *n* of the RNA to ensure comparability between RNAs of different size.

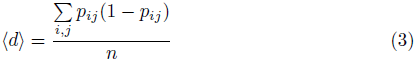

The distance between the ensemble and the reference structure, 〈*d*(*S*)〉, is a quantitative measure for the agreement of the predicted ensemble with the accepted target structure. In contrast to the *Pairing Probability Score*, which focuses on the predicted pairing probabilities for base pairs present in the target structure, the probabilities for all possible basepairs are taken into account. Since the incorporation of experimental data into prediction algorithms tends to prefer structures being in accordance with the determined structural features, the ensemble distance can be used to quantify the expected shift of the whole ensemble towards the reference structure.

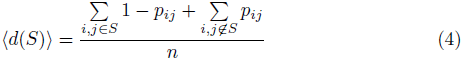

### 3 Benchmark Data

The test set created by Hajdin *et al.* (2013) was used for benchmarking the accuracy of secondary structure predictions including SHAPE data (http://www.chem.unc.edu/rna/data-files/ShapeKnots_DATA.zip). The test set contains 24 sequences with their corresponding SHAPE data sets and reference structures, which are required to rate the prediction results. The reference structures were derived from X-ray crystallography experiments or predicted by comparative sequence analysis. As shown in Figure 1, the benchmark shows a high diversity regarding the length and prediction *Sensitivity* of the involved RNAs.

**Figure 1:**
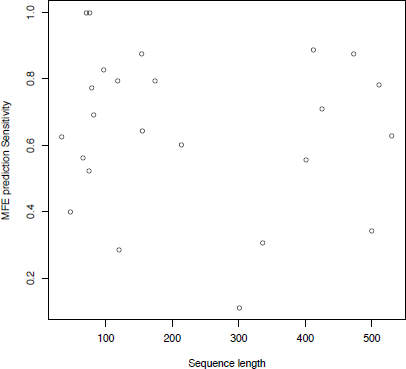
Length and MFE prediction *Sensitivity* of RNAs used for benchmarking.

The mentioned test set has been designed for benchmarking the prediction of secondary structures containing pseudoknots, which scales at *O*(*n*^6^). In contrast to secondary structure prediction without pseudoknots, which scales at *O*(*n*^3^), the computational effort is growing much faster with growing sequence lengths. As a result the longest RNA sequences in the test set have a length of about 500 nucleotides. Nevertheless, the benchmark also contains structures without pseudoknots. A perfect prediction for structures containing pseudoknots is not possible, since the ViennaRNA Package does not support pseudoknots. Therefore, we applied a simple optimization strategy that removes pseudo-knots from the reference structures while keeping the number of base pairs maximal.

All three implemented methods depend on a set of carefully adjusted parameters. While we use the published default parameters for the methods of Deigan *et al.* (2009) (*m* = 1.8, *b* = −0.6) and Zarringhalam *et al.* (2012) (*β* = 0.8), we performed an exhaustive evaluation of the parameter space for the method presented by Washietl *et al.* (2012), see Figure 2. Based on this data, we selected the following default parameter combinations for this approach: *τ/σ* = 2.0, minimizer tolerance *є_m_* = 0.001, initial step size of the minimizer method *s_m_* = 0.01. Furthermore, our implementation of the method by Washietl *et al.* (2012) in the program RNApvmin defaults to an estimation of the gradient by drawing 1000 sample structures from the Boltzmann ensemble. This not only considerably speeds up the optimization routines, but also enables their application to rugged landscapes where an exact gradient approach could easily trap the optimization process in a local minimum. The benchmark results for all three methods that correspond to their beforementioned default parameters are listed in Table 1. **RNApvmin results need to be updated in the following table**

**Table 1:**
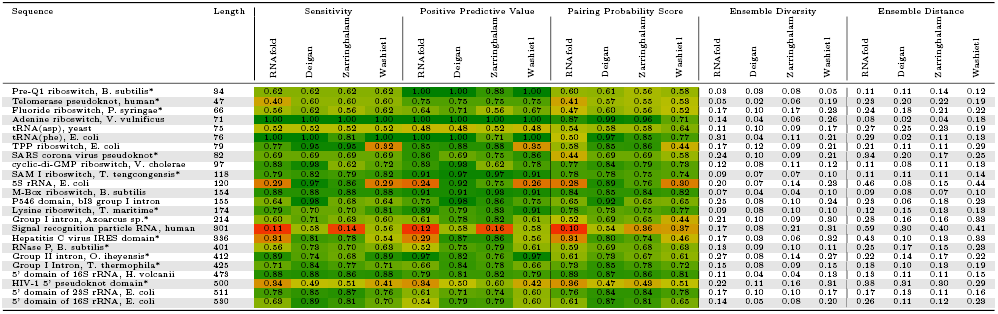
*Sensitivity, Positive Predictive Value, Pairing Probability Score, Ensemble Diversity* and distance between ensemble and reference structure for the secondary structure predictions created by thermodynamic based folding and SHAPE directed folding according to Deigan et al., Zarringhalam et al. and Washietl et al.

**Figure 2:**
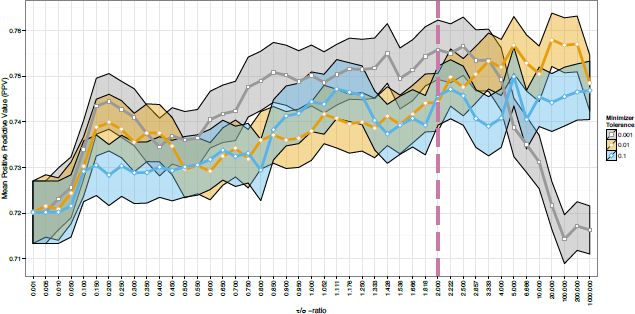
Parameter space evaluation for the method of Washietl *et al.* (2012). Plotted are the mean positive predictive values (PPV) for the entire benchmark data set using different parameter settings. For the sake of clarity, only three different values for the minimizer tolerance *є_m_*, namely 0.1, 0.01, and 0.001, are depicted, while for each of them a large range of *τ/σ*-ratios is used. The polygon surrounding each line of mean values indicates the standard deviation of PPVs within the entire set of predictions for the corresponding parameter setting. The dashed, purple, vertical line highlights the *τ/σ*-ratio used as default value for `RNApvmin`.

### 4 Theophylline sensing riboswitch

As an additional, particular example for comparing the three presented methods we selected the artificially designed RNA switch *theo-P-is10*, taken from Qi *et al.* (2012). This RNA consists of a theophylline sensing aptamer part followed by a ncRNA expression platform. The switching principle follows a regular ON-switch behavior, where under sufficiently high concentrations of theophylline, the aptamer part of the structure is thermodynamically favored, and the downstream located ncRNA part of the sequence folds into its active state. On the other hand, in the absence of theophylline, the expression platform misfolds into an inactive state.

In the original design *theo-P-is10* forms a pseudoknot interaction between the aptamer stem and the expression platform in the inactive state. However, by using `RNAfold` we explicitly exclude pseudoknots, which is also true for almost all other secondary structure prediction programs available. Nevertheless, the data that comes with the work of Qi *et al.* (2012) provides a rich source of interesting SHAPE probing data, since it consists of normalized reactivities from two experiments: (i) the RNA folds in theophylline free solution, and (ii) the RNA folds in the presence of 0.5 mM theophylline, respectively. Since the designed pseudoknot is only present in the inactive state of the RNA switch, i.e. in theophylline-free solution, we do not emphasize too much on the correctness of the structure prediction in this case.

To compare the different variants of guided RNA secondary structure prediction through SHAPE reactivity data incorporation for *theo-P-is10*, we computed the ground state structures, and base pair probabilities for the two corresponding experimental data sets. Instead of using two dotplots for comparison, we create differential `RNAbow` plots (Aalberts and Jannen, 2013) to visualize the difference in base pair probability predictions. Here, a differential `RNAbow` plot consists of two sets of arcs located on the upper and lower half of the horizontally aligned nucleotide sequence, showing the predictions for both experiments, respectively. The strength/width of the arcs represents the pairing probability (thicker lines mean higher probability), whereas arcs are colored with an intensity corresponding to the absolute value of difference in predictions (red in the upper half, blue in the lower half) only, if pairing probability is higher when compared to the other experiment. For better visualization we restrict the `RNAbow` plots to pairing probabilities of 0.1 and above.

Figure 3 outlines the two ground state structures of the designed RNA switch together with their corresponding pairing probabilities in form of a bowplot. Results of the predictions using the method of Deigan *et al.* (2009) with default parameters, the parameters used in Qi *et al.* (2012), the method of Zarringhalam *et al.* (2012) with default parameters, and the method of Washietl *et al.* (2012), are shown in Figures 4, 5, 6, and 7, respectively. It can be easily seen that the method of Deigan *et al.* (2009) using default parameters clearly misses the proposed ground state structures and essentially yields results as obtained by `RNAfold` without incorporation of SHAPE reactivities. On the other hand, using the two parameters *m* = 3.4, and *b* = −0.5, both SHAPE reactivity data sets yield high probabilities for the aptamer pocket and the functional ncRNA part, although only in presence of theophylline the aptamer pocket is fully formed. Using the method of Zarringhalam *et al.* (2012) both predicted ground state structures again correspond to the active conformation of the designed RNA switch. However, the proposed pseudoknot interaction between the two hairpin loops of the inactive state becomes visible in the base pair plot. This effect is even more pronounced when using the method of Washietl *et al.* (2012). Here, the pairing probabilities for the pseudoknot interaction are much higher in the absence of theophylline, whereas the probabilities of the base pairs involved in the formation of the aptamer pocket and the ncRNA part are increased in the presence of theophylline. Nevertheless, both ground state structures are virtually identical and represent the active conformation.

**Figure 3:**
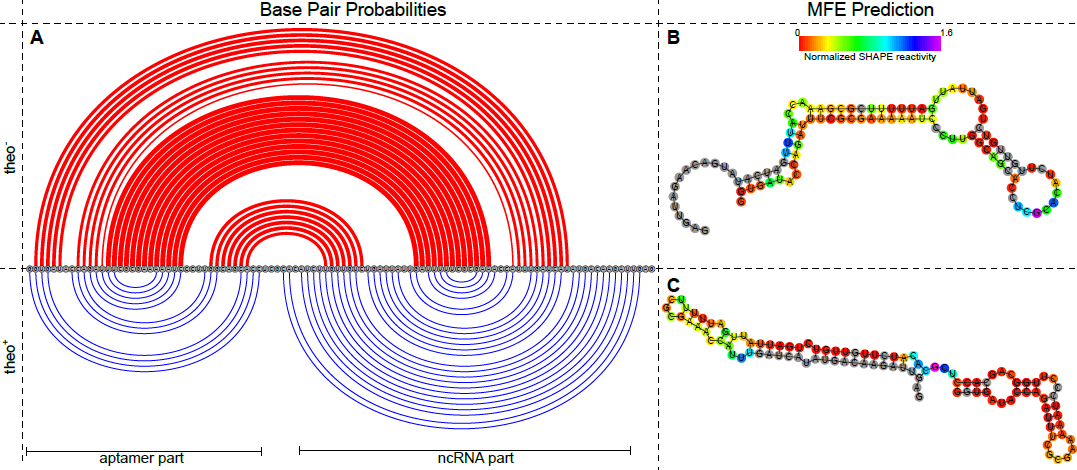
The model system of the designed theophylline Riboswitch (Qi *et al.*, 2012). **A** Base pair probabilities of the two proposed secondary structure states as computed by `RNAfold` depicted by an `RNAbow` diagram. Similar to the differential `RNAbow` plot, base pair probabilities are depicted by the strength/width of the arcs. However, the upper half (red arcs) shows the base pairs of the MFE structure (supposedly OFF state), and the lower half (blue arcs) the ON-state, respectively, instead of the difference between two distinct predictions. Secondary structure drawings of the MFE structure and the ON-state with a color encoding of their respective SHAPE reactivity as measured by Qi *et al.* (2012) are shown in **B**, and **C**, respectively.

**Figure 4:**
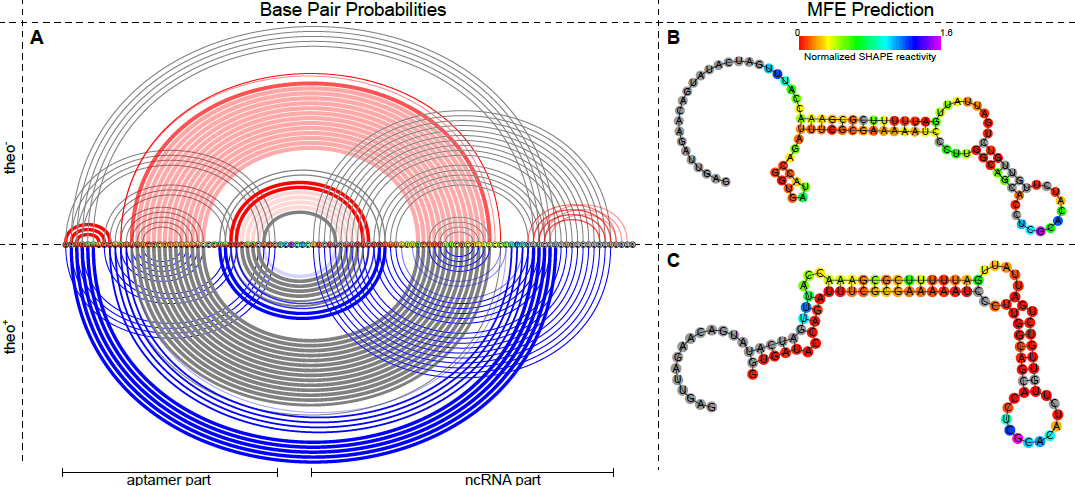
Guided secondary structure prediction with RNAfold using the method of Deigan *et al.* (2009) with default parameters (*m* = 1.8 and *b* = −0.6). **A** Differential `RNAbow` diagram of the base pair probabilities (see text). SHAPE reactivity data for the theophylline-free probing experiment (upper half), and a concentration of 0.5 mM theophylline (lower half) are taken from Qi *et al.* (2012). Although the (short-range) aptamer part and ncRNA part of *theo-P-is10* show relatively high pairing probabilities, especially in the presence of theophylline (lower half, blue arcs), long-range interactions dominate the structure ensemble in both cases. The two secondary structure plots **B**, and **C** show the predicted MFE structures for the theophylline-free system, and under presence of theophylline, respectively. Colored nucleotides show the corresponding SHAPE reactivity data, whereas grey circles indicate the absence of probing data.

**Figure 5:**
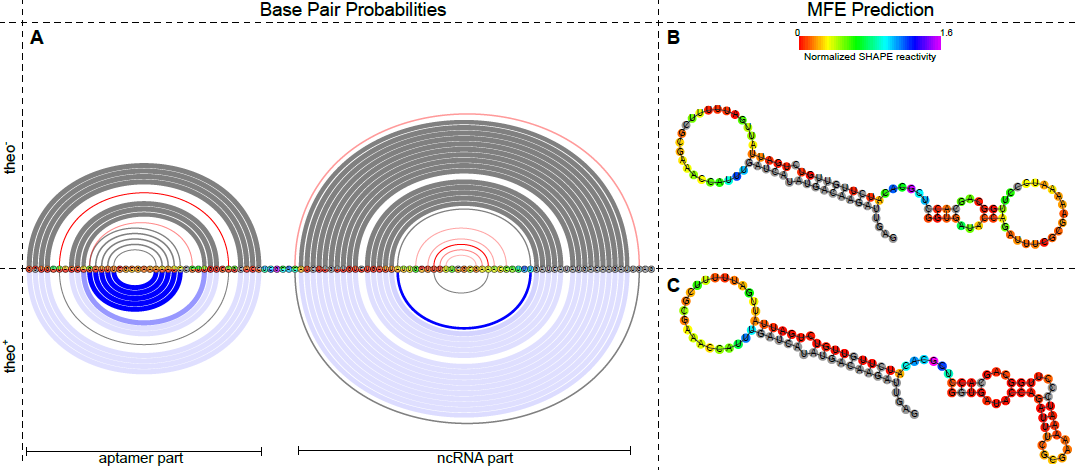
Guided secondary structure prediction with RNAfold using the method of Deigan *et al.* (2009) with parameters taken from Qi *et al.* (2012) (*m* = 3.4 and *b* = −0.5). **A** Differential `RNAbow` diagram of the base pair probabilities. SHAPE reactivity data for the theophylline-free probing experiment (upper half), and a concentration of 0.5 mM theophylline (lower half) are taken from Qi *et al.* (2012). Here, both predictions show a shift of the structure ensemble towards the functional ON-state of the RNA switch. Except for the innermost stem of the aptamer part of the designed RNA, pairing probabilities of the (dominant) functional state are only slightly higher in the presence of theophylline (light blue). The two secondary structure plots **B**, and **C** show the predicted MFE structures for the theophylline-free system, and under presence of theophylline, respectively. Colored nucleotides show the corresponding SHAPE reactivity data, whereas grey circles indicate the absence of probing data.

**Figure 6:**
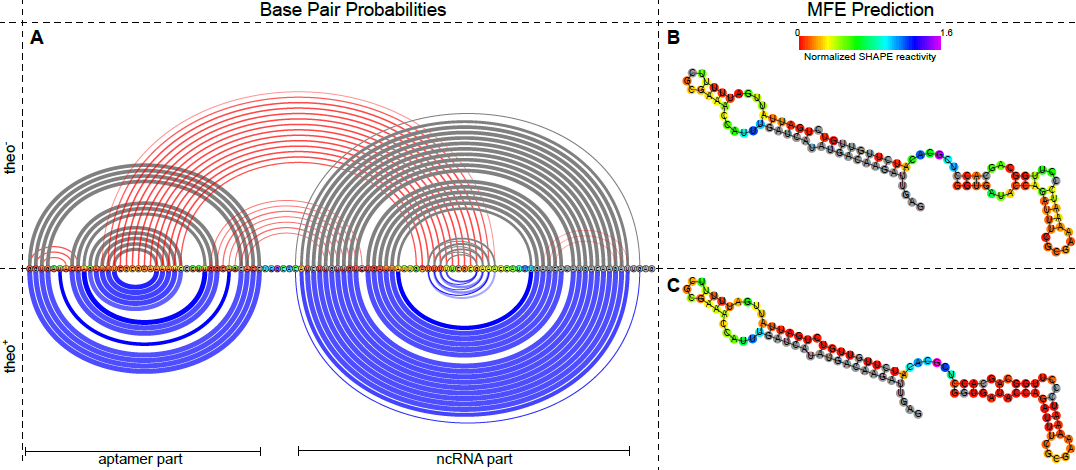
Guided secondary structure prediction with RNAfold using the method of Zarringhalam *et al.* (2012) with default parameters. **A** Differential `RNAbow` diagram of the computed base pair probabilities. SHAPE reactivity data for the theophylline-free probing experiment (upper half), and a concentration of 0.5 mM theophylline (lower half) are taken from Qi *et al.* (2012). Both data sets result in high probabilities for the aptamer, and ncRNA part. However, in the presence of theophylline, the possible long-range interaction between the two complementary hairpin loop sequences of these parts (upper half, large red arcs) falls below the probability threshold of 0.1 used for the construction of the `RNAbow` plot. This goes along with a high shift of the ensemble towards the ON-state of the riboswitch (lower half, blue arcs). The two secondary structure plots **B**, and **C** show the predicted MFE structures for the theophylline-free system, and under presence of theophylline, respectively. Colored nucleotides show the corresponding SHAPE reactivity data, whereas grey circles indicate the absence of probing data. Both SHAPE reactivity data sets result in a MFE prediction of the ON-state, with the only difference of an additional A-U base pair in the ligand binding pocket of the aptamer part under the presence of theophylline.

**Figure 7:**
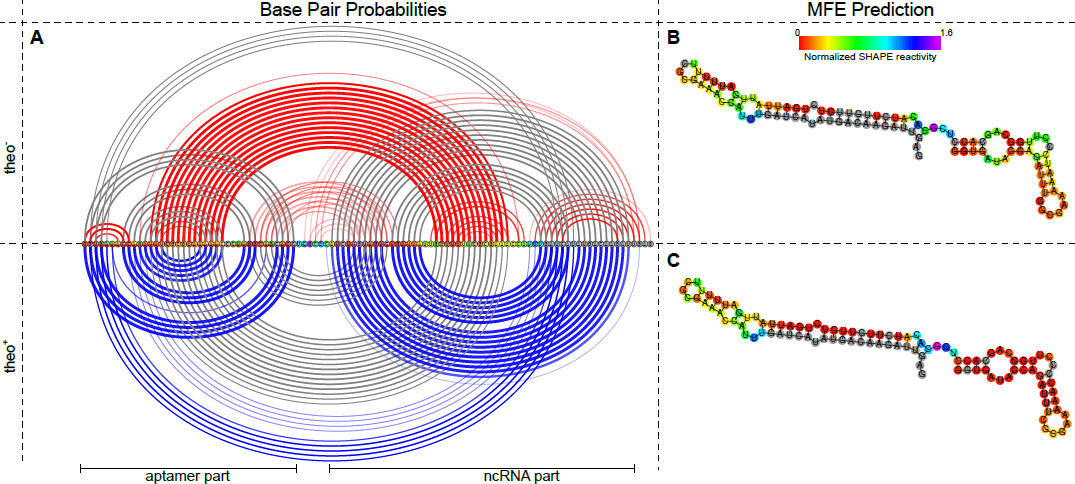
Guided secondary structure prediction with RNAfold using the method of Washietl *et al.* (2012) with default parameters. Here, the two provided SHAPE reactivity data sets were used to compute a position-wise perturbation vector for the usage in guided secondary structure prediction. **A** Differential `RNAbow` diagram of the computed base pair probabilities. SHAPE reactivity data for the theophylline-free probing experiment (upper half), and a concentration of 0.5 mM theophylline (lower half) are taken from Qi *et al.* (2012). The two different modes (OFF/ON state) are clearly visible in the differential probabilities. While under theophylline-free condition (upper half) a long range interaction between the two complementary hairpin loop parts of the aptamer and the ncRNA show rather high probability (red arcs), this is not the case under the presence of theophylline (lower half). Here, the base pairs of the aptamer part as well as the ncRNA part of the RNA switch exhibit distinctly high pairing probability (dark blue arcs). The two secondary structure plots **B**, and **C** show the predicted MFE structures for the theophylline-free system, and under presence of theophylline, respectively. Colored nucleotides show the corresponding SHAPE reactivity data, whereas grey circles indicate the absence of probing data. Although the pairing probabilities as shown in **A** would suggest otherwise, both predicted ground state structures represent the ON-state of the RNA switch.

In contrast to the above methods, the implementation of so-called *soft-constraints* in the ViennaRNA Package 2.2 (published elsewhere) also allows for a direct inclusion of binding free energies of the ligand to the aptamer pocket. For this purpose, the ensemble of structures is modified such that all structures that exhibit the aptamer pocket receive an additional stabilizing free energy of *E_s_* = −9.22 kcal/mol, according to the dissociation constant of *K_d_* = 0.32*μM* taken from Jenison *et al.* (1994). The resulting constrained secondary structure prediction is shown in Figure 8. Here, the shift towards the functional ligand binding state of the RNA switch under presence of theophylline is clearly visible in the base pair probabilities.

**Figure 8:**
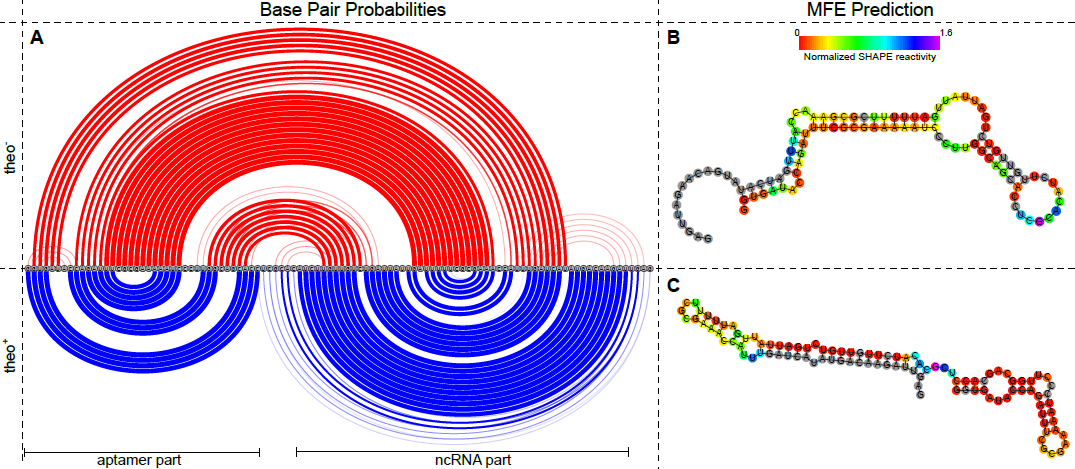
Direct incorporation of theophylline binding free energy using the *soft constraints* feature of the ViennaRNA Package 2.2. Here, structures in the lower part of the figure (*theo^+^*) receive an additional stabilizing free energy of *E_s_* = −9.22 kcal/mol if they exhibit the aptamer pocket. This soft constraint is derived from the dissociation constant *K_d_* = 0.32*μM* taken from Jenison *et al.* (1994)−9.22. The upper part (*theo*^−^) consists of data obtained by `RNAfold` with default parameters. **A** Differential `RNAbow` diagram of the computed base pair probabilities. The two secondary structure plots **B**, and **C** show the predicted MFE structures for the theophylline-free system, and under presence of theophylline, respectively. For comparison with the corresponding experimentally measured SHAPE reactivity data, nucleotides are colored according to this data. Grey circles indicate the absence of probing data. The ON and OFF state of the RNA switch are clearly visible in both, the ground state and the base pair probability predictions.

### 5 Mapping SHAPE reactivities to pairing probabilities

While the approach of Deigan et al. directly converts SHAPE reactivities to pseudo energies, the methods of Zarringhalam et al. and Washietl et al. both require experimentally determined pairing probabilities as input data. However, converting raw reactivity values to pairing probabilities is not a trivial task and both approaches use different methods to calculate pairing probabilities based on given SHAPE reactivities. While Washietl et al. used a simple cutoff approach to distinguish between paired and unpaired positions, Zarringhalam et al. used a more sophisticated method where the normalization is carried out in a stepwise linear fashion (See table 2).

**Table 2:** Linear mapping classes used to convert SHAPE reactivities to probabilities for being unpaired according to Zarringhalam et al.

| SHAPE reactivity | Probability for being unpaired |
| --- | --- |
| <0.25 | 0.00 - 0.35 |
| 0.25 - 0.30 | 0.35 - 0.55 |
| 0.30 - 0.70 | 0.55 - 0.85 |
| >0.70 | 0.85 - 1.00 |

In this benchmark a common method was used to compute the required pairing probabilities based on the experimentally determined SHAPE reactivities. The application RNAplfold was used to predict the pairing probabilities for all sequences of the benchmark. The predicted pairing probabilities of all nucleotides were then compared with the determined SHAPE reactivities. The dataset containing about 4500 observations showed a significant correlation between the logarithm of the SHAPE reactivity and the probability for a certain nucleotide to be unpaired. However, as shown in figure 9, the experimental signal shows a high variation and high reactivities can also be observed for paired nucleotides, and vice versa. Nevertheless, a linear model is suitable for converting the logarithm of the SHAPE reactivity to the probability for being unpaired. 

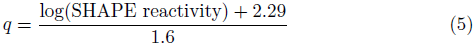

**Figure 9:**
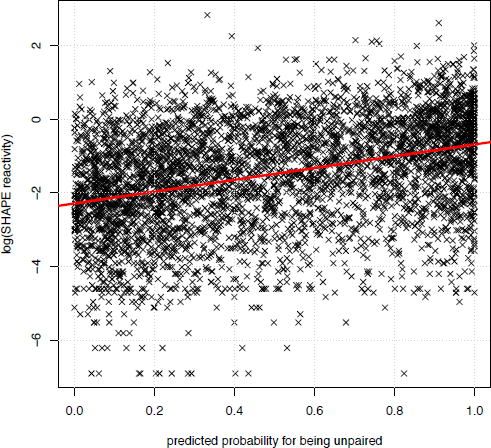
Relation of measured SHAPE reactivities to predicted probabilities for being unpaired

Since the equation above may also lead to results lower than 0 and higher than 1, all results exceeding those limits are mapped to the corresponding boundary value.

### 6 Runtime

The impact of the incorporation of additional soft constraints onto the required amount of computational time was benchmarked for the whole dataset using a workstation (Intel Core 2 Quad 2.83 GHz, GCC 4.8.2). The runtime for the folding recursion was averaged over 10 runs. As shown in figure 10, the incorporation of additional constraints results in a slight increase of the required computational time. However, the effect is less distinct for the approach of Deigan et al. which may be explained by the fact that in contrast to the other approaches, which apply pseudo energies for every paired/unpaired nucleotide, the free energy is only adapted when evaluating stacked pairs.

**Figure 10:**
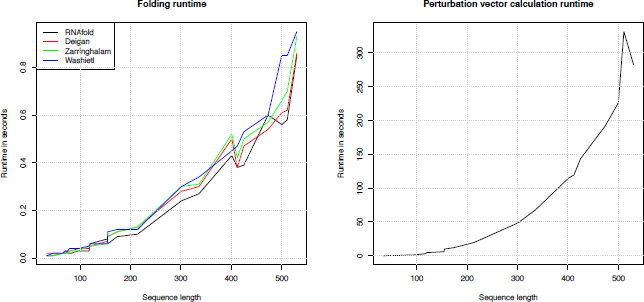
Runtime for predicting the MFE structure and the partition function using various approaches and runtime required to calculate a perturbation vector with exact gradient evaluation

**Figure 11:**
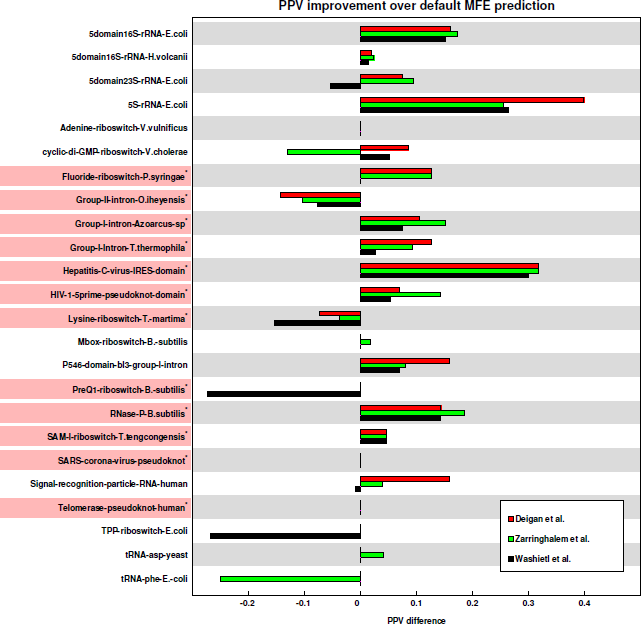
Change of the positive predictive value (PPV) of the minimum free energy structure due to constrained folding compared to unconstrained folding. Reference structures that contain pseudoknots are marked with by a superscripted asterisk and light-red background. The poor performance of the Washietl *et al.* (2012) method in the case of *E.coli TPP riboswitch* is caused by an inconsistency between SHAPE reactivity and proposed reference structure. On the other hand, the *PreQ1-riboswitch in B.subtilis* is an extremely short, heavily pseudoknotted example where a prediction using the Washietl *et al.* (2012) method yields almost no base pairs.

**Figure 12:**
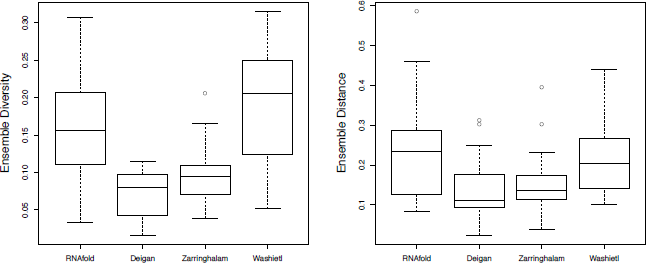
*Ensemble Diversity* and distance between ensemble and reference structure for unconstrained and constrained structure predictions

The overall runtime for the prediction according to Washietl et al. can be separated into two phases. First, a perturbation vector is calculated by numerically minimizing an objective function. This step requires most of the computational resources since the exact evaluation of the gradient at various points of the minimization algorithm scales at *0*(*N*^4^). However, the evaluation can be done much faster when the gradient is estimated from a number of sampled sequences. Second, the calculated perturbation vector is used to constrain the secondary structure prediction.

